# The role of accelerated growth plate fusion in the absence of SOCS2 on osteoarthritis vulnerability

**DOI:** 10.1101/2021.05.13.444074

**Authors:** Hasmik J. Samvelyan, Carmen Huesa, Lucy Cui Lin, Colin Farquharson, Katherine A. Staines

## Abstract

Osteoarthritis is the most prevalent systemic musculoskeletal disorder characterised by articular cartilage degeneration and subchondral bone (SCB) sclerosis. Here we sought to examine the contribution of accelerated growth to osteoarthritis development using a murine model of excessive longitudinal growth. Suppressor of cytokine signalling 2 (SOCS2) is a negative regulator of growth hormone (GH) signalling, thus mice deficient in SOCS2 (*Socs2*^-/-^) display accelerated bone growth. We examined vulnerability of *Socs2*^-/-^ mice to osteoarthritis following surgical induction of disease (destabilisation of the medial meniscus (DMM)), and with ageing, by histology and micro-CT. We observed significant increase in number (WT DMM: 532±56; WT sham: 495±45; KO DMM: 169±49; KO sham: 187±56; P<0.01) and density (WT DMM: 2.2±0.9; WT sham: 1.2±0.5; KO DMM: 13.0±0.5; KO sham: 14.4±0.7) of growth plate bridges in *Socs2*^-/-^ in comparison to wild-type (WT). Histological examination of WT and *Socs2*^-/-^ knees revealed articular cartilage damage with DMM in comparison to sham (WT DMM: 3.4±0.4; WT sham: 0.3±0.05 (P<0.05); KO DMM: 3.2±0.8; KO sham: 0.8±0.3). Articular cartilage lesion severity scores (mean and maximum) were similar in WT and *Socs2*^-/-^ mice with either DMM, or with ageing. Micro-CT analysis revealed significant decreases in SCB thickness, epiphyseal trabecular number and thickness in the medial compartment of *Socs2*^-/-^, in comparison to WT (P<0.001). DMM had no effect on the SCB thickness in comparison to sham in either genotype. Together these data suggest that enhanced GH signalling through SOCS2 deletion accelerates growth plate fusion, however this has no effect on osteoarthritis vulnerability in this model.

**Summary statement:** Deletion of SOCS2 results in accelerated growth plate fusion, however this has no effect on osteoarthritis vulnerability.

## Introduction

Osteoarthritis is the most prevalent systemic musculoskeletal disorder characterised by degeneration of joint articular cartilage, osteophyte formation, subchondral bone plate thickening, synovial proliferation and inflammation. Osteoarthritis has a multifactorial aetiology including ageing, trauma, obesity and heredity. Further, a complex interplay of major molecules and signalling pathways play indispensable roles in osteoarthritis development (Chang et al., 2019; Echtermeyer et al., 2009; Glasson et al., 2005; Saito et al., 2010; Wang et al., 2013). Despite this, effective disease-modifying treatments are currently limited.

Endochondral ossification is an essential process for longitudinal bone growth. It requires hypertrophic differentiation of chondrocytes characterised by secretion of type X collagen (COL10A1), matrix metalloproteinase-13 (MMP-13) and vascular endothelial growth factor (VEGF), followed by the subsequent degradation and conversion of the cartilage matrix into highly vascularised bone tissue (Kronenberg, 2003; Nagao et al., 2017). Osteoarthritis is widely accepted to involve the reversion of chondrocyte behaviour to an earlier developmental-like phenotype, which could drive the disease process. Indeed, re-expression of the type IIA procollagen, a spliced variant of the type II collagen gene (COL2A1) normally expressed in chondroprogenitor cells, in adult osteoarthritic articular chondrocytes indicates reversion of these cells to early developmental-like phenotype (Aigner et al., 1999). Consistent with this, we have previously shown the abnormal deployment of a transient chondrocyte phenotype in the joints of a STR/ort mouse (Staines et al., 2016), a murine model for spontaneous osteoarthritis (Samvelyan et al., 2020). Further, we revealed accelerated long bone growth, a wider zone of growth plate proliferative chondrocytes, and widespread COL10A1 and MMP-13 expression beyond the expected hypertrophic zone distribution in these mice, which may underpin their osteoarthritis onset (Staines et al., 2016). However, the precise contribution of accelerated growth to osteoarthritis development remains unclear.

Suppressor of cytokine signalling 2 (SOCS2), one of the members of the suppressor of cytokine signalling family glycoproteins, is implicated in cancer and disorders of immune system and central nervous system (Cramer et al., 2019; Keating and Nicholson, 2019; Letellier and Haan, 2016). SOCS2 has been shown as a primary intracellular suppressor of growth hormone signalling pathway, thus mice deficient in SOCS2 (*Socs2*^-/-^) display an excessive growth phenotype (Metcalf et al., 2000). Here we use this mouse model of accelerated bone growth to understand the association between aberrant growth dynamics and osteoarthritis development. To achieve this, we examined the vulnerability of *Socs2*^-/-^ mice to osteoarthritis following surgical induction of disease (destabilisation of the medial meniscus (DMM)), and with ageing, by histology and micro computed tomography (micro-CT).

## Results

### Socs2^*-/-*^ mice exhibit accelerated growth plate fusion

In accordance with the known effects of increased growth hormone (GH) signalling on the skeleton, we observed a significant increase in *Socs2*^-/-^ body weight in comparison to WT controls in ageing (Socs2^-/-^ 51.9g ± 1.4; WT 32.0g ± 0.5g; P<0.001), and throughout the 8-week DMM experiment (Table 1; P<0.001). We next sought to examine growth plate fusion in these mice. We observed a significant increase in the number of growth plate bridges in *Socs2*^-/-^ mice in comparison to WT mice at both ages examined (Fig. 1). Specifically, we saw a significant increase in growth plate bridge number (Fig. 1A & B; WT DMM: 532 ± 56; WT sham: 495 ± 45; KO DMM: 169 ± 49; KO sham: 187 ± 56; P<0.01) and density (Fig. 1A & C; (WT DMM: 2.2 ± 0.9; WT sham: 1.2 ± 0.5; KO DMM: 13.0 ± 0.5; KO sham: 14.4 ± 0.7; P<0.01) in *Socs2*^-/-^ sham and DMM tibiae (16 weeks of age), in comparison to WT sham and DMM tibiae (P<0.01). No differences in growth plate bridges were observed with osteoarthritis intervention in either genotype. Concurrent with this, in aged mice (around 1 year), a significant increase in growth plate bridge number (Fig. 1D & E) and density (Fig. 1D & F) was observed in *Socs2*^-/-^ mice in comparison to WT mice (WT: 585 ± 70; KO: 1659 ± 91; densities – WT: 8.6 ± 2.2; KO: 21.3 ± 1.4 P<0.001).

**Table 1:**
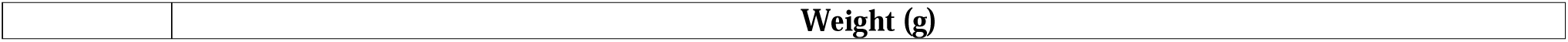
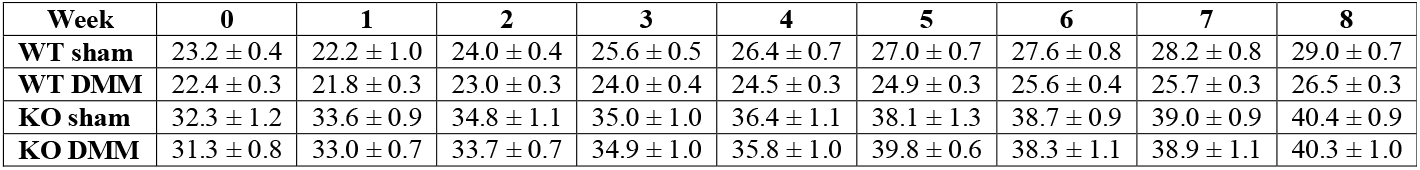
Weights of wild type (WT) and Socs2^-/-^ (KO) mice during 8-week post-DMM experimental timeline.

| **Week** | **0** | **1** | **2** | **3** | **4** | **5** | **6** | **7** | **8** |
| --- | --- | --- | --- | --- | --- | --- | --- | --- | --- |
| **WT sham** | 23.2 ± 0.4 | 22.2 ± 1.0 | 24.0 ± 0.4 | 25.6 ± 0.5 | 26.4 ± 0.7 | 27.0 ± 0.7 | 27.6 ± 0.8 | 28.2 ± 0.8 | 29.0 ± 0.7 |
| **WT DMM** | 22.4 ± 0.3 | 21.8 ± 0.3 | 23.0 ± 0.3 | 24.0 ± 0.4 | 24.5 ± 0.3 | 24.9 ± 0.3 | 25.6 ± 0.4 | 25.7 ± 0.3 | 26.5 ± 0.3 |
| **KO sham** | 32.3 ± 1.2 | 33.6 ± 0.9 | 34.8 ± 1.1 | 35.0 ± 1.0 | 36.4 ± 1.1 | 38.1 ± 1.3 | 38.7 ± 0.9 | 39.0 ± 0.9 | 40.4 ± 0.9 |
| **KO DMM** | 31.3 ± 0.8 | 33.0 ± 0.7 | 33.7 ± 0.7 | 34.9 ± 1.0 | 35.8 ± 1.0 | 39.8 ± 0.6 | 38.3 ± 1.1 | 38.9 ± 1.1 | 40.3 ± 1.0 |

**Figure 1.**
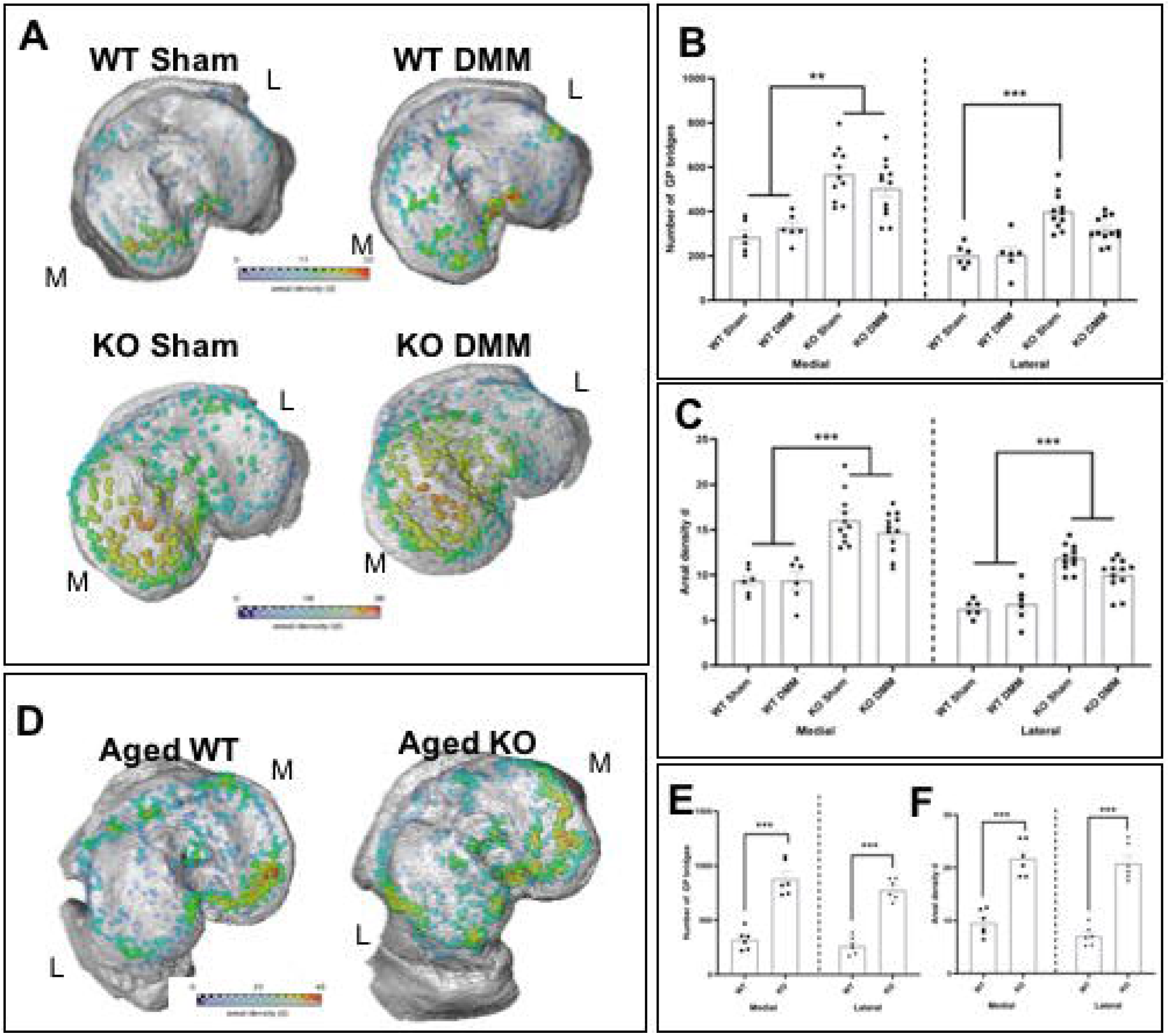
Socs2 deficient mice exhibit accelerated growth plate fusion mechanisms. **(A)** Location and areal densities of bridges across the growth plate projected on the medial (M) and lateral (L) tibial joint surface in WT and Socs2^-/-^ (KO) DMM and sham operated joints. **(B)** Number of bridges. **(C)** Areal density (d) of bridges defined as the number of bridges per 256 mm x 256 mm window. **(D)** Location and areal densities of bridges across the growth plate projected on the medial (M) and lateral (L) tibial joint surface in aged WT and Socs2^-/-^ (KO) mice. **(E)** Number of bridges **(F)** Areal density (d) of bridges defined as the number of bridges per 256 mm x 256 mm window. Data are presented as mean ± SEM and showing individual animals. Statistical test performed: Two-way ANOVA with Bonferroni adjustments for multiple comparisons within each joint compartment. * *p*<0.05 ** *p*<0.01 *** *p*<0.001

### Socs2 deletion does not exacerbate the development of osteoarthritis in a DMM model

Assessment of cartilage damage in the medial tibia of WT mice revealed an increased articular cartilage OARSI score with DMM in comparison to sham (P<0.05; Fig. 2A, C & D). However, no significant differences in the articular cartilage mean and maximum OARSI severity scores were observed between WT and *Socs2*^-/-^ mice (Fig. 2C & 2D). Similarly, whilst osteophytes were observed in both WT and *Socs2*^-/-^ DMM mice, there was no difference between genotypes (Fig. 2B). Micro-CT analysis of the subchondral bone plate revealed a significantly thinner subchondral bone plate in the medial compartment of *Socs2*^-/-^ knee joints, in comparison to WT knee joints (Fig. 3A; SCB Th. WT: 0.15mm ± 0.003; KO: 0.11mm ± 0.003; P<0.001). DMM had no effect on the subchondral bone plate thickness in comparison to sham in either WT or Socs2^-/-^ knee joints (Fig. 3A). Similarly, no differences were observed in the subchondral bone % BV/TV between genotypes or osteoarthritis interventions (Fig. 3B). Micro-CT analysis of the epiphyseal trabecular bone also revealed no effect of DMM on % BV/TV (Fig. 3C), however we observed decrease in trabecular number (Fig. 3D; Tb. N. WT: 12.3mm^-1^ ± 0.2; KO: 10.6mm^-1^ ± 0.2; P<0.001) and increase in trabecular thickness (Fig. 3E; Tb. Th. WT: 0.06mm ± 0.001; KO: 0.07mm ± 0.002; P<0.05) in *Socs2*^-/-^ knee joints in comparison to WT knee joints. DMM had no effect on the epiphyseal bone parameters in comparison to sham in either WT or *Socs2*^-/-^ knee joints (Fig. 3C - F).

**Figure 2.**
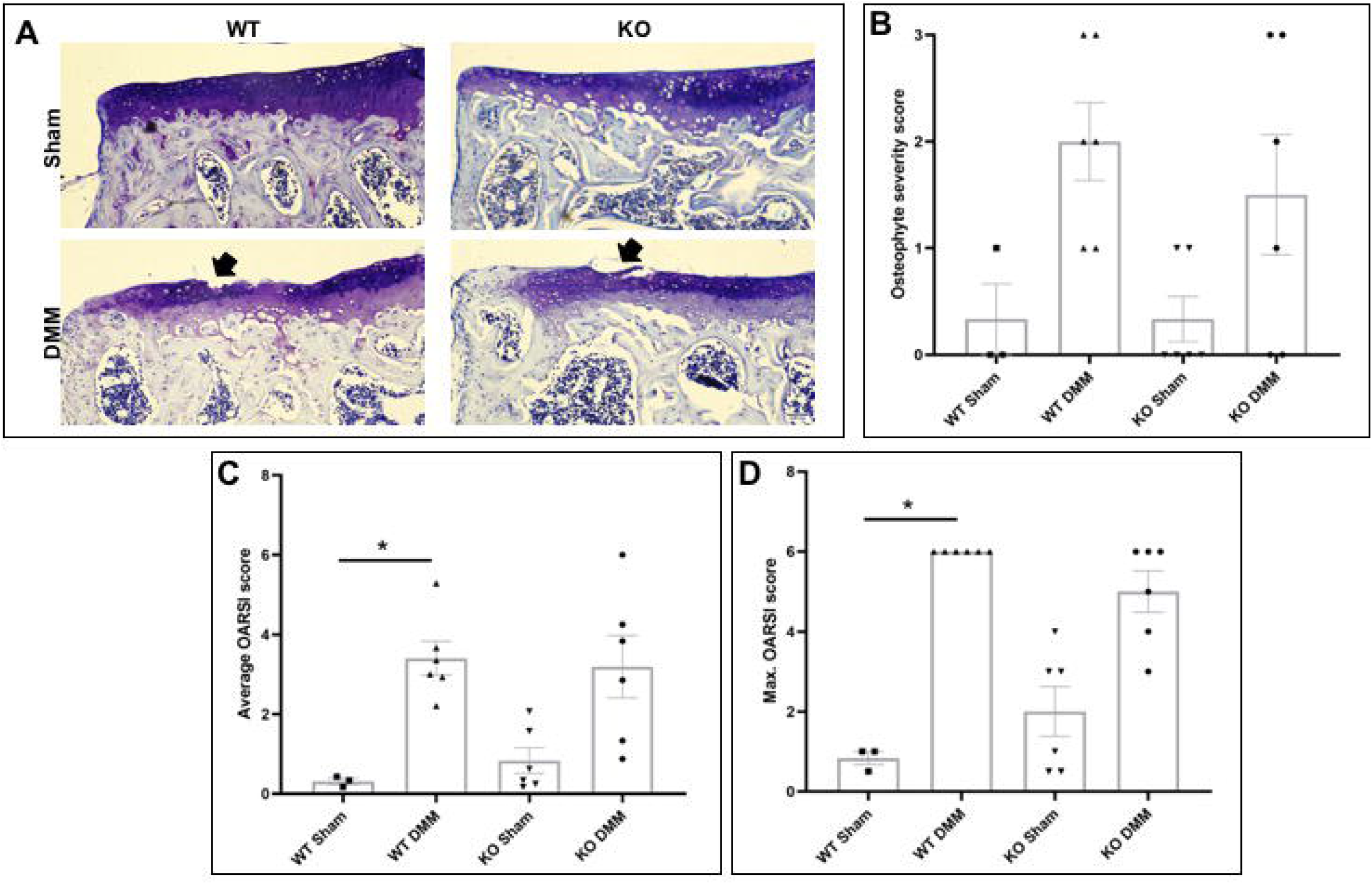
Deletion of Socs2 does not prevent osteoarthritic articular cartilage lesions in a surgical model of osteoarthritis. **(A)** Toluidine blue stained sections of the knee joint of WT and Socs2^-/-^ (KO) mice showing development of articular cartilage lesions in the medial tibia. **(B)** Osteophyte severity score **(C)** Mean articular cartilage damage OARSI score and **(D)** Maximum articular cartilage damage OARSI score in WT sham (n = 3), WT DMM (n = 6), KO sham (n = 6) and KO DMM (n = 6). Scale bar = 0.05mm. Data are presented as mean ± SEM and showing individual animals. Statistical test performed: Kruskal-Wallis one-way ANOVA.

**Figure 3.**
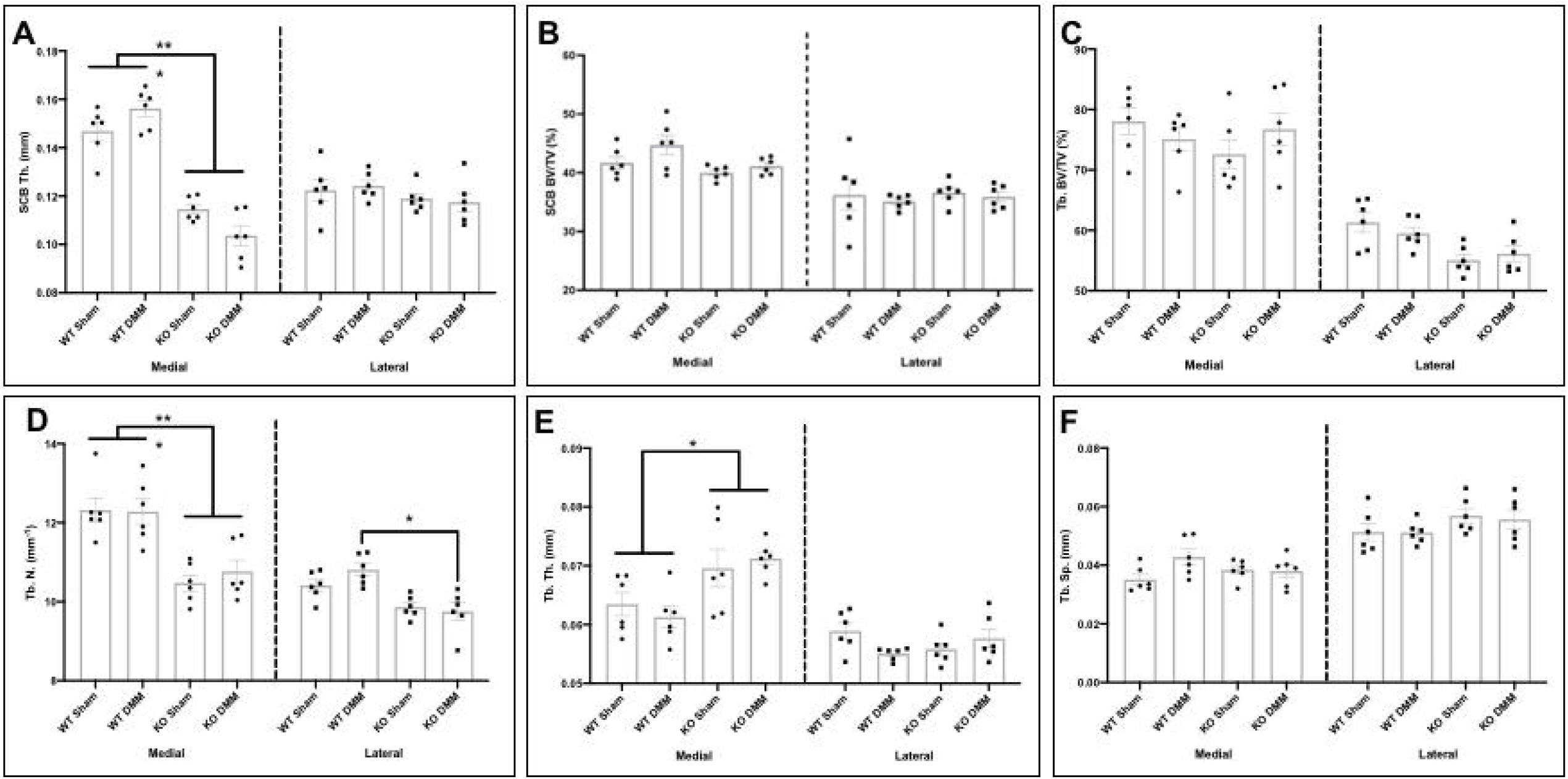
Socs2 deficient mice exhibit decreased subchondral bone plate and trabecular number, irrespective of DMM surgically induced osteoar thritis. Micro-CT analysis of the medial and lateral tibial **(A)** Subchondral bone plate (SCB) thickness (mm) **(B)** SCB bone volume/tissue volume (BV/TV; %) **(C)** Epiphyseal trabecular BV/TV (%) **(D)** Epiphyseal trabecular number (mm^-1^) **(E)** Epiphyseal trabecular thickness (mm) **(F)** Epiphyseal trabecular separation (mm) in WT and Socs2^-/-^ (KO) mice with DMM or sham surgery (n=6/group). Data are presented as mean ± SEM and showing individual animal data. Statistical test performed: Two-way ANOVA with Bonferroni adjustments for multiple comparisons. *P<0.05; ***P<0.001.

### Socs2 deletion has no effect on joint ageing

To examine the effects of *Socs2* deletion in aged joints, histological examination revealed no differences in the articular cartilage lesion mean and maximum severity scores between aged WT and Socs2^-/-^ mice in any of the joint compartments (Figs. 4A, B & 4C). Similar to our previous analysis of DMM treated WT and *Socs2*^-/-^ mice (Fig. 3), micro-CT analysis of the subchondral bone revealed thinner subchondral bone plate in the medial compartment of *Socs2*^-/-^ knee joints, in comparison to WT knee joints (Fig. 5A; SCB Th. WT: 0.2mm ± 0.005; KO: 0.1mm ± 0.004; P<0.001). No differences were observed in the subchondral bone % BV/TV between genotypes (Fig. 5B), however the epiphyseal trabecular BV/TV was decreased in *Socs2*^-/-^ knee joints (Fig. 5C; Tb. BV/TV WT: 73.5% ± 1.5; KO: 64.6% ± 2.2; P<0.01). No significant differences were observed in trabecular number (Fig. 5D), trabecular thickness (Fig. 5E) or trabecular separation (Fig. 5F) between aged WT and Socs2^-/-^ mice.

**Figure 4.**
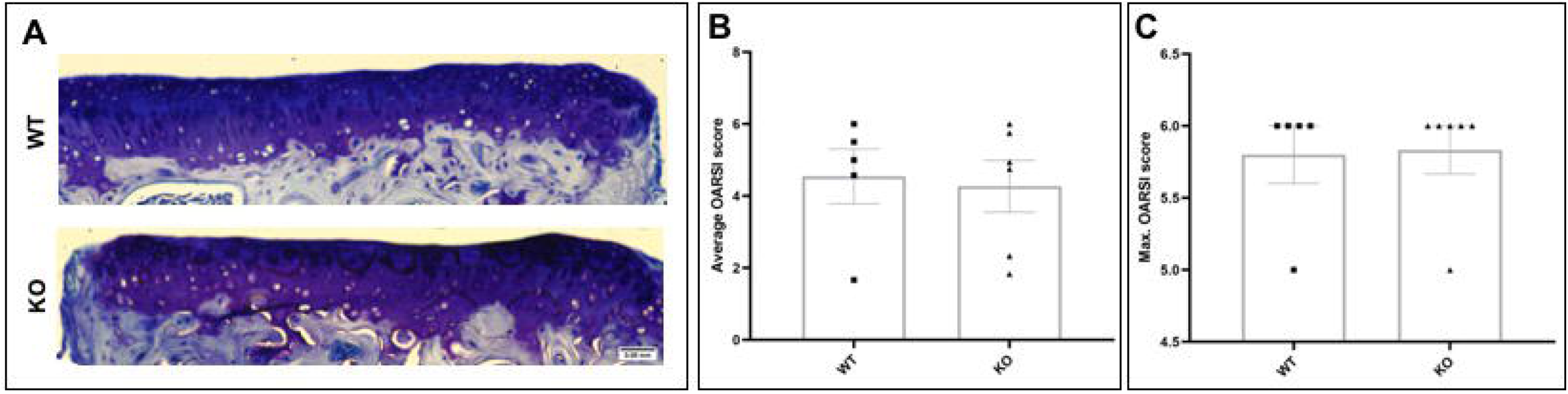
Deletion of Socs2 does not prevent osteoarthritic articular cartilage lesions with ageing. **(A)** Toluidine blue stained sections of the knee joint of WT and Socs2^-/-^ (KO) mice showing development of articular cartilage lesions in the medial tibia. **(B)** Mean articular cartilage damage OARSI score across the knee joint and **(C)** Maximum articular cartilage damage OARSI score between WT (n=5) and KO (n=6) mice with ageing, in the medial tibia of the knee joint. Scale bar = 0.05mm. Data are presented as mean ± SEM and showing individual animal data. Statistical test performed: Mann-Whitney U test.

**Figure 5.**
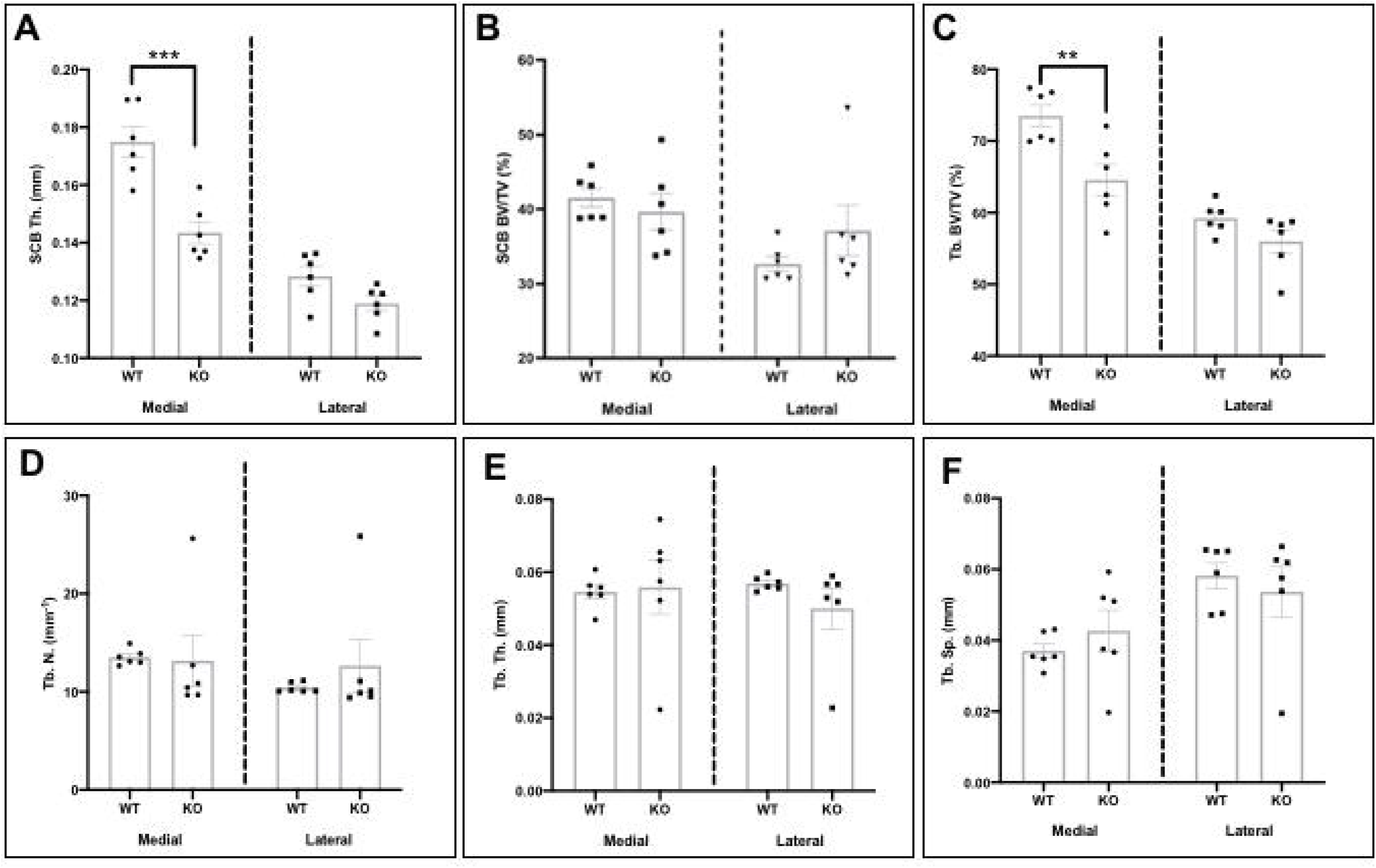
Aged Socs2 deficient mice exhibit decreased subchondral bone plate thickness. Micro-CT analysis of the medial and lateral tibial **(A)** Subchondral bone plate (SCB) thickness (mm) **(B)** SCB bone volume/tissue volume (BV/TV; %) **(C)** Epiphyseal trabecular BV/TV (%) **(D)** Epiphyseal trabecular number (mm^-1^) **(E)** Epiphyseal trabecular thickness (mm) **(F)** Epiphyseal trabecular separation (mm) in aged WT and Socs2^-/-^ (KO) mice (n=6/group). Data are presented as mean ± SEM and showing individual animals. Statistical test performed: Two-way ANOVA with Bonferroni adjustments for multiple comparisons. *P<0.05; ***P<0.001.

## Discussion

Here we sought to examine whether altered growth plate dynamics underpin osteoarthritis through examination of osteoarthritis vulnerability in a murine model of accelerated growth (*Socs2*^-/-^). We describe accelerated growth plate fusion in these mice, consistent with their known overgrowth phenotype. However, we found no effect of a surgical intervention or ageing on the articular cartilage or subchondral bone phenotype in these mice. This suggests that in this murine model, aberrant growth dynamics are not associated with vulnerability to osteoarthritis development.

It is well established that in osteoarthritis, chondrocytes in the articular cartilage adopt a more transient phenotype, similar to that seen in the growth plate (Pitsillides and Beier, 2011). This raises the question as to whether a greater understanding of the discordant chondrocyte phenotypes may inform on mechanisms underpinning osteoarthritis, and strategies for treatment. Our previous work has shown that in a spontaneous model of osteoarthritis (STR/Ort mouse), there is an association between aberrant growth plate dynamics and osteoarthritis development (Staines et al., 2016). Specifically, we revealed STR/Ort mice show an overgrowth phenotype with enriched growth plate bridging, which was associated with articular cartilage lesions at 18-20 weeks of age (Staines et al., 2016). Further, our previous work on the MRC National Survey of Health and Development revealed modest associations between greater gains in height in childhood, indicative of accelerated growth, and decreased odds of knee osteoarthritis at 53 years (Staines et al., 2020). Together this suggests that the growth rate may play a role in the development of osteoarthritis, although what that role is has yet to be fully defined.

SOCS2 is a negative regulator of GH signalling, via inhibition of the Janus kinase/signal transducers and activators of transcription (JAK/STAT) pathway (Pass et al., 2009). Thus, mice deficient in *Socs2*^-/-^ display an excessive growth phenotype (Metcalf et al., 2000). Characterisation of their growth phenotype has revealed increased bone growth rates, growth plate widths, and chondrocyte proliferation in *Socs2*^-/-^ 6□week□old mice compared to age-matched wild-type mice (Pass et al., 2012). *Socs2*^-/-^ mice have normal serum levels of GH and insulin-like growth factor -1 (IGF-1), and their longitudinal growth phenotype is due to local effects of the GH/IGF-1 axis on the growth plate (Macrae et al., 2009). Consistent with this, we revealed increased numbers and densities of growth plate bridges in *Socs2*^-/-^ mice, in comparison to WT mice. Growth plate bridges form as the growth slows and undergoes progressive narrowing. They are also known to form upon growth plate injury, thought to be through an intramembranous ossification mechanism (Xian et al., 2004). However, whether growth plate fusion occurs prior to or after the cessation of growth is of significant controversy in the field and has been somewhat overlooked (Parfitt, 2002).

We have previously shown using finite element modelling growth plate bridging to increase stress dissipation in the subchondral bone region of the joint (Madi et al., 2019; Staines et al., 2018). It is therefore surprising that the increased growth plate bridging observed in our *Socs2*^-/-^ mice here had reduced subchondral bone plate and trabecular parameters. Previous studies have examined the *Socs2*^-/-^ bone phenotype with contradictory results. Our previous work has shown that *Socs2*^-/-^ mice have increased bone mass, trabecular number and trabecular thickness (Dobie et al., 2018; Macrae et al., 2009). However, others have shown the absence of SOCS2 to induce losses in cortical and trabecular bone mineral density (Lorentzon et al., 2005). These results are not consistent with the expected augmented GH/IGF-1 axis, but are consistent with our findings here and highlight the complexity of this pathway and the need for further studies to elucidate the precise role of SOCS2 signalling in bone homeostasis.

Increased GH but reduced IGF-1 concentrations are present in synovial fluid of patients with osteoarthritis (Denko et al., 1996). However, in rodents deficiency of GH and IGF-1 causes an increased severity of osteoarthritic articular cartilage lesions (Ekenstedt et al., 2006). Further, it has previously been shown that *Socs2* mRNA levels are decreased in chondrocytes from osteoarthritic femoral heads (de Andres et al., 2011). Together these data suggest a role for the GH/IGF-1/SOCS2 pathway in the pathology of osteoarthritis. However, our results presented here indicate that there is no effect of SOCS2 deficiency on osteoarthritis vulnerability, and thus highlighting the need to better understand GH/IGF-1 signalling in the aetiology of osteoarthritis. This may be due to the normal serum levels of GH/IGF-1 observed in *Socs2*^-/-^ mice (Macrae et al., 2009).

Together, our data show that deletion of SOCS2 leads to accelerated growth plate fusion, but this has no effect on osteoarthritis vulnerability in a surgical model of osteoarthritis. Future studies will determine whether this lack of vulnerability is specific to this model of accelerated longitudinal growth, or whether this is characteristic of osteoarthritis in general.

## Methods

### Animals

*Socs2*^□*/*□^ mice on a C57/BL6 genetic background were generated as previously described (Dobie et al., 2018). For genotyping, tail biopsied DNA was analysed by PCR for *Socs2* WT (Forward: TGTTTGACTGAGCTCGCGC, Reverse: CAACTTTAGTGTCTTGGATCT) or the neocassette (*Socs2* ^*-/-*^; Forward: ACCCTGCACACTCTCGTTTTG, Reverse: CCTCGACTAAACACATGTAAAGC). Mice were kept in polypropylene cages, with light/dark 12-h cycles, at 21 ± 2°C, and fed *ad libitum* with maintenance diet (Special Diet Services, Witham, UK). Ageing studies were completed in 12-13-month-old *Socs2*^□*/*□^ (n=6) and C57/BL6 wild-type (WT; n=6) male mice (Charles River). Analyses were conducted blindly to minimise the effects of subjective bias. All experimental protocols were approved by Roslin Institute’s Animal Users Committee and the animals were maintained in accordance with UK Home Office guidelines for the care and use of laboratory animals.

### Destabilisation of the medial meniscus (DMM)

Osteoarthritis was induced in 8-week old *Socs2*^□*/*□^ (n=6) and C57/BL6 WT (n=6) male mice (Charles River) by surgically induced DMM under isoflurane-induced anaesthesia. Following transection of the medial meniscotibial ligament to destabilise the medial meniscus, the left knee joint capsule and skin were closed and anaesthesia reversed. Sham□operated joints were used as controls. After 8 weeks, knee joints were dissected, fixed in 4% paraformaldehyde for 24 hours at 4^°^C, and then stored in 70% ethanol.

### Micro-CT analysis

Scans of the right knee joint were performed with an 1172 X-Ray microtomograph (Skyscan, Belgium) to evaluate the subchondral bone. High-resolution scans with voxel size of 5 µm were acquired (50 kV, 200µA, 0.5 mm aluminium filter, 0.6° rotation angle). The projection images were reconstructed using NRecon software version 1.6.9.4 (Skyscan, Belgium). Each dataset was rotated in DataViewer (Skysan, Belgium) to ensure similar orientation and alignment for analysis. Hand-drawn regions of interests (ROI) of the subchondral trabecular bone for each femur and tibia lateral/medial compartments was first achieved. Subchondral bone ROIs was subsequently selected for each compartment. Analysis of subchondral bone plate thickness and the epiphyseal trabecular bone was achieved using 3D algorithms in CTAn (Skyscan, Belgium) to provide: subchondral bone plate thickness (SCB Th.; mm); subchondral bone plate bone volume/tissue volume (SCB BV/TV; %); epiphyseal trabecular bone volume/tissue volume (Tb. BV/TV; %); trabecular number (Tb. N.; mm^-1^); trabecular thickness (Tb. Th.; mm) and trabecular separation (Tb. Sp.; mm).

### Growth plate bridging analysis

Growth plate bridging analysis was conducted using a 3D micro-CT quantification method as previously described (Madi et al., 2019; Staines et al., 2018). Briefly, micro-CT scans of the tibiae were segmented using a region-growing algorithm within the Avizo® (V8.0, VSG, Burlington, VT, USA) software. The central points of all bony bridges were identified and projected on the tibial joint surface. The distribution of the areal number density of bridges (N, the number of bridges per 256 µm × 256 µm window) is then calculated and superimposed on the tibial joint surface (each bridge has a colour that represents the areal number density at the bridge location).

### Histological analysis

Murine left knee joints were decalcified, wax-embedded and 6μm coronal sections cut. For assessment of osteoarthritis severity, multiple sections (>5/slide) from 120μm intervals across the whole joint were stained with Toluidine blue (0.4% in 0.1 M acetate buffer, pH 4). Articular cartilage damage was assessed in the medial tibia using the well-established OARSI grading scale, with scores averaged (Glasson et al., 2010). Osteophytes were also scored where 0 = none; 1 = formation of cartilage-like tissue; 2 = increase in cartilaginous matrix; 3 = endochondral ossification (Nagira et al., 2020). Scoring was conducted blindly with by two observers.

### Statistical analysis

All analyses were performed with GraphPad Prism software 6.0f version (GraphPad Inc, La Jolla, CA, USA). The results were presented as the mean ± standard error of the mean (SEM). Normal distribution of data was assessed using the Shapiro-Wilk normality test. For the growth plate bridging and micro-CT analysis, two-way ANOVA with Bonferroni adjustments for multiple comparisons were used. For articular cartilage damage, the Kruskal-Wallis one-way ANOVA was used for DMM studies, and the Mann-Whitney U test for ageing studies. The significance was set at P<0.05.

## Acknowledgements

The authors would like to thank Dr Blandine Poulet of University of Liverpool, Institute of Ageing and Chronic Disease for her guidance in experimental techniques and insightful discussions throughout preparation.

## Data availability

Data are available upon reasonable request to the corresponding author.

## Author contributions

Conception and design of the study: KAS, CF, HJS. Acquisition of data: HJS, KAS, CH, LCL. Drafting the manuscript: HJS, KAS. Revising the manuscript and final approval, and agreement to be accountable for all aspects of the work: all authors.

